# Repeat Exposure to Hypercapnic Seawater Modifies Performance and Oxidative Status in a Tolerant Burrowing Clam

**DOI:** 10.1101/2020.08.03.234955

**Authors:** Samuel J. Gurr, Shelly A. Trigg, Brent Vadopalas, Steven B. Roberts, Hollie M. Putnam

## Abstract

Moderate oxidative stress is a hypothesized driver of enhanced stress tolerance and lifespan. Whereas thermal stress, irradiance, and dietary restriction show evidence of dose-dependent benefits for many taxa, stress acclimation remains understudied in marine invertebrates, despite being threatened by climate change stressors such as ocean acidification. To test for life-stage and stress-intensity dependence in eliciting enhanced tolerance under subsequent stress encounters, we initially conditioned pediveliger Pacific geoduck (*Panopea generosa*) larvae to (i) ambient and moderately elevated *p*CO_2_ (920 μatm and 2800 μatm, respectively) for 110 days, (ii) secondarily applied a 7-day exposure to ambient, moderate, and severely elevated *p*CO_2_ (750 μatm, 2800 μatm, and 4900 μatm, respectively), followed by 7 days in ambient conditions, and (iii) implemented a modified-reciprocal 7-day tertiary exposure to ambient (970 μatm) and moderate *p*CO_2_ (3000 μatm). Initial conditioning to moderate *p*CO_2_ stress followed by secondary and tertiary exposure to severe and moderate *p*CO_2_ stress increased respiration rate, organic biomass, and shell size suggesting a stress-intensity-dependent effect on energetics. Additionally, stress-acclimated clams had lower antioxidant capacity compared to clams under initial ambient conditions, supporting the hypothesis that stress over postlarval-to-juvenile development affects oxidative status later in life. We posit two subcellular mechanisms underpinning stress-intensity-dependent effects on mitochondrial pathways and energy partitioning: i) stress-induced attenuation of mitochondrial function and ii) adaptive mitochondrial shift under moderate stress. Time series and stress intensity-specific approaches can reveal life-stages and magnitudes of exposure, respectively, that may elicit beneficial phenotypic variation.

**Summary statement:** Hypercapnic conditions during postlarval development improves physiological performance and oxidative status. This novel investigation of adaptive stress acclimation highlights the plasticity of bioenergetic and subcellular responses in *Panopea generosa*.

## 1. Introduction

Ocean acidification (OA), or the decrease of oceanic pH, carbonate ion concentration, and aragonite saturation (*Ω*) due to elevated atmospheric partial pressures (*p*CO_2_), poses a global threat with magnified intensity in coastal marine systems (Cai et al., 2011). Marine molluscs are particularly susceptible to OA, with negative physiological impacts in aerobic performance (Navarro et al., 2013), calcification, growth and development (Waldbusser et al., 2015), acid/base regulation (Michaelidis et al., 2005), and energy-consuming processes (i.e. protein synthesis; Pan et al., 2015).

Principles in ectotherm physiology (i.e. oxygen capacity-limited thermal tolerance: Pörtner, 2012; energy-limited tolerance to stress: Sokolova, 2013) describe aerobic performance “windows” under ‘optimum’, ‘pejus’ (moderate), and ‘pessimum’ (severe) environmental ranges defined by cellular and physiological modifications affecting energy homeostasis (Sokolova et al., 2012). The conserved defense proteome, or cellular stress response (CSR), is the hallmark of cellular protection but comes at an energetic cost (Kültz, 2005). Whereas the CSR is unsustainable if harmful conditions exacerbate or persist (Sokolova et al., 2012), episodic or sublethal stress encounters can induce adaptive phenotypic variation (Tanner and Dowd, 2019). A growing body of research suggests moderate or intermittent stress (e.g. caloric restriction, irradiance, thermal stress, oxygen deprivation, etc.) can elicit experience-mediated resilience for a variety of taxa (i.e. fruit fly, coral, fish, zebra finch, mice) increasing CSR, fitness, and compensatory/anticipatory responses under subsequent stress exposures (Brown et al., 2002; Costantini et al., 2012; Jonsson and Jonsson, 2014; Visser et al., 2018; Zhang et al., 2018). Further, early-life development presents a sensitive stage to elicit adaptive phenotypic adjustments (Fawcett and Frankenhuis, 2015) prompting investigation of environmental stress acclimation under a rapidly changing environment.

Dose-dependent stress response, or conditioning-hormesis (Calabrese et al., 2007), explains how stress priming can enhance tolerance limitations under subsequent encounters to similar or higher levels of stress intensity later in life (Costantini, 2014). Oxidative stress presents a common source of conditioning-hormesis (Costantini, 2014) and is a hypothesized driver of longevity (Ristow and Schmeisser, 2014; Wojtczyk-Miaskowska and Schlichtholz, 2018). For example, early-life exposure to moderate oxidative stress in the Caribbean fruit fly *Anastrepha suspensa* and zebra finch *Taeniopygia guttata* decreases cellular damage and increases proteomic defense, energy assimilation, and survival under a subsequent stress encounter during adulthood (Costantini et al., 2012; Visser et al., 2018). Oxidative stress, or an over-production of reactive oxygen species (ROS; superoxide, hydrogen peroxide, hydroxyl radical) primarily from mitochondrial oxidative phosphorylation, causes macromolecular damage. In marine invertebrates, oxidative stress can intensify under environmental stressors such as hypoxia and emersion (Abele et al., 2008), hyposalinity (Tomanek et al., 2012), thermal stress (An and Choi, 2010), pollutants and contaminants (Livingstone, 2001), and OA (Tomanek et al., 2011; Matoo et al., 2013). Protein families that are involved in the CSR, function in signaling, avoidance, and mediation of oxidative damage. Specifically, antioxidant proteins (i.e. superoxide dismutase, catalase, glutathione peroxidase, etc.) are widely conserved across phyla to scavenge ROS and ameliorate redox status at the expense of energy homeostasis (Kültz, 2005). Adaptive cellular defense against oxidative damage is thought to have an important evolutionary role in the longevity of the ocean quahog *Arctica islandica* (lifespan > 400 years) due to a lifestyle of metabolic dormancy (when burrowed) and aerobic recovery (Abele et al., 2008). Further, Ivanina and Sokolova (Ivanina and Sokolova, 2016) found hypoxia-tolerant marine bivalves show anticipatory and compensatory upregulation of antioxidant proteins to mitigate oxidative bursts under hypoxia-reoxygenation. Such adaptive responses have yet to be explored under hypercapnic conditions to identify species tolerant to OA stress. Although bivalves are known to exhibit *p*CO_2_-induced oxidative damage and upregulated CSR (Tomanek et al., 2011; Matoo et al., 2013), studies have yet to investigate ROS-mediated activity in a hormetic framework (repeated dose exposures).

Pacific geoduck (*Panopea generosa*) are burrowing clams of ecological (Goodwin and Pease, 1987) and economic importance (Shamshak and King, 2015) and are a great candidate for investigating a hormetic framework for generation of stress-acclimated phenotypes. Juvenile geoduck have shown positive carryover effects after exposures under high *p*CO_2_/low *Ω* conditions including compensatory respiration rates and shell growth (Gurr et al., 2020). In contrast, larval performance is negatively impacted under OA exposure (Timmins-Schiffman et al., 2020). The postlarval life stage presents an ecologically relevant and less susceptible window to investigate effects of *p*CO_2_ stress acclimation. ‘Settlement’ in bivalves is a developmental transition from free-swimming larvae in an oxygen-saturated water column to an increasingly sedentary or burrowed life in the benthos (Goodwin and Pease, 1989) where stratification, bacterial carbon mineralization, and reduced buffering capacity drives down aragonite saturation and oxygen (Cai et al., 2011). To investigate thresholds of hormesis and potential for beneficial stress acclimation, we investigated the effects of *p*CO_2_ exposures of different intensity and at different time points in a repeated reciprocal fashion, on the physiological and subcellular phenotypes of juvenile Pacific geoduck.

## 2. Materials and Methods

### 2.1 Experimental setup

Larval Pacific geoduck were reared from gametes at the Jamestown Point Whitney Shellfish Hatchery (Brinnon, WA) following standard shellfish aquaculture industry practices, using bag-filtered (5μm) and UV sterilized seawater pumped from offshore (27.5 m depth) in Dabob Bay (WA, USA). Ambient seawater temperature, salinity, pH, and *p*CO_2_ was 16-18 °C, 29 ppt, 7.7-7.8 pH, and ~800-950 μatm, respectively. Larvae reached settlement competency, characterized by a protruding foot and larval shell length >300 μm, at 30 days post-fertilization. Approximately 15,000 larvae were randomly placed into each of eight 10-L trays (Heath/Tecna) containing a thin layer of sand to simulate the natural environment and enable metamorphosis from veliger larvae to pediveliger larvae, and subsequently to the burrowing and sessile juvenile stage.

### 2.2 Acclimation from pediveligers to juveniles

#### Initial Exposure (110 days)

Pediveligers were placed into ambient and elevated *p*CO_2_ conditions (921 ± 41 μatm and 2870 ± 65 μatm; Table 1; Fig. 1) for an initial exposure during the transition from pediveliger to the burrowing juvenile stage (*N*=4 trays treatment^−1^; *N*=1.5×10^4^ pediveligers tray^−1^). Seawater flowed into 250-L head tanks at a rate of 0.1 L min^−1^ and replicate trays were gravity-fed from the head tanks. At the end of the initial exposure, respiration rate and shell growth was measured for 20 randomly selected juveniles from each of the eight trays as described below. Additionally, six animals from each tray were frozen in liquid nitrogen and stored at −80 °C for molecular analyses.

**Table 1.**
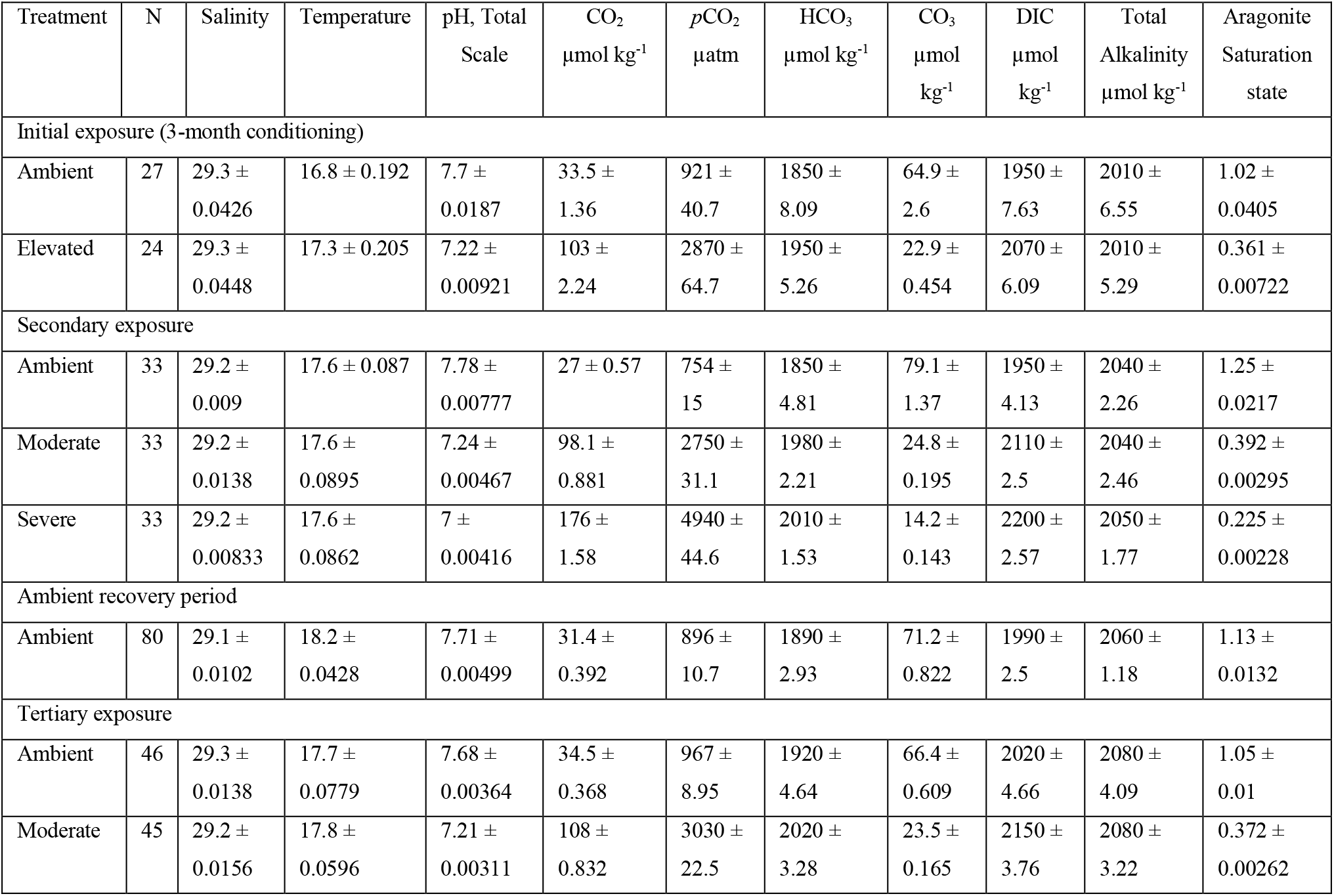
Seawater carbonate chemistry. pH, salinity, and temperature measured with handheld probes and total alkalinity (via Gran titration) measured with 60 mL from trays and tanks during the 110-day acclimation period (weekly) and during the 21-day experiment, respectively. Seawater carbonate chemistry (CO_2_, *p*CO_2_, HCO_3_^−^, CO_3_^−2^, DIC, aragonite saturation state) was calculated with the seacarb R package (Gattuso et al., 2018).

| Treatment | N | Salinity | Temperature | pH, Total Scale | $\text{CO}_2$<br>$\mu\text{mol kg}^{-1}$ | $p\text{CO}_2$<br>$\mu\text{atm}$ | $\text{HCO}_3^-$<br>$\mu\text{mol kg}^{-1}$ | $\text{CO}_3^{2-}$<br>$\mu\text{mol kg}^{-1}$ | DIC<br>$\mu\text{mol kg}^{-1}$ | Total Alkalinity<br>$\mu\text{mol kg}^{-1}$ | Aragonite Saturation state |
| --- | --- | --- | --- | --- | --- | --- | --- | --- | --- | --- | --- |
| Initial exposure (3-month conditioning) |  |  |  |  |  |  |  |  |  |  |  |
| Ambient | 27 | $29.3 \pm 0.0426$ | $16.8 \pm 0.192$ | $7.7 \pm 0.0187$ | $33.5 \pm 1.36$ | $921 \pm 40.7$ | $1850 \pm 8.09$ | $64.9 \pm 2.6$ | $1950 \pm 7.63$ | $2010 \pm 6.55$ | $1.02 \pm 0.0405$ |
| Elevated | 24 | $29.3 \pm 0.0448$ | $17.3 \pm 0.205$ | $7.22 \pm 0.00921$ | $103 \pm 2.24$ | $2870 \pm 64.7$ | $1950 \pm 5.26$ | $22.9 \pm 0.454$ | $2070 \pm 6.09$ | $2010 \pm 5.29$ | $0.361 \pm 0.00722$ |
| Secondary exposure |  |  |  |  |  |  |  |  |  |  |  |
| Ambient | 33 | $29.2 \pm 0.009$ | $17.6 \pm 0.087$ | $7.78 \pm 0.00777$ | $27 \pm 0.57$ | $754 \pm 15$ | $1850 \pm 4.81$ | $79.1 \pm 1.37$ | $1950 \pm 4.13$ | $2040 \pm 2.26$ | $1.25 \pm 0.0217$ |
| Moderate | 33 | $29.2 \pm 0.0138$ | $17.6 \pm 0.0895$ | $7.24 \pm 0.00467$ | $98.1 \pm 0.881$ | $2750 \pm 31.1$ | $1980 \pm 2.21$ | $24.8 \pm 0.195$ | $2110 \pm 2.5$ | $2040 \pm 2.46$ | $0.392 \pm 0.00295$ |
| Severe | 33 | $29.2 \pm 0.00833$ | $17.6 \pm 0.0862$ | $7 \pm 0.00416$ | $176 \pm 1.58$ | $4940 \pm 44.6$ | $2010 \pm 1.53$ | $14.2 \pm 0.143$ | $2200 \pm 2.57$ | $2050 \pm 1.77$ | $0.225 \pm 0.00228$ |
| Ambient recovery period |  |  |  |  |  |  |  |  |  |  |  |
| Ambient | 80 | $29.1 \pm 0.0102$ | $18.2 \pm 0.0428$ | $7.71 \pm 0.00499$ | $31.4 \pm 0.392$ | $896 \pm 10.7$ | $1890 \pm 2.93$ | $71.2 \pm 0.822$ | $1990 \pm 2.5$ | $2060 \pm 1.18$ | $1.13 \pm 0.0132$ |
| Tertiary exposure |  |  |  |  |  |  |  |  |  |  |  |
| Ambient | 46 | $29.3 \pm 0.0138$ | $17.7 \pm 0.0779$ | $7.68 \pm 0.00364$ | $34.5 \pm 0.368$ | $967 \pm 8.95$ | $1920 \pm 4.64$ | $66.4 \pm 0.609$ | $2020 \pm 4.66$ | $2080 \pm 4.09$ | $1.05 \pm 0.01$ |
| Moderate | 45 | $29.2 \pm 0.0156$ | $17.8 \pm 0.0596$ | $7.21 \pm 0.00311$ | $108 \pm 0.832$ | $3030 \pm 22.5$ | $2020 \pm 3.28$ | $23.5 \pm 0.165$ | $2150 \pm 3.76$ | $2080 \pm 3.22$ | $0.372 \pm 0.00262$ |

**Fig. 1.**
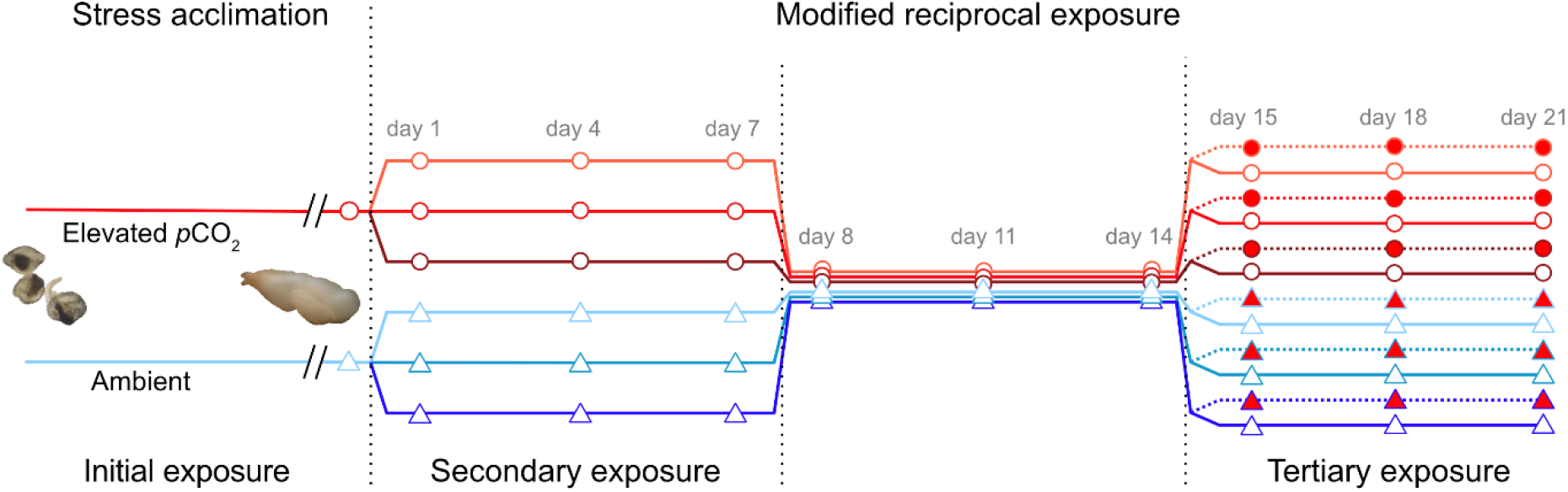
Schematic of the experimental design. Line color and shape refers to the *p*CO_2_ treatments during the initial 110-day acclimation period (blue triangles, ambient *p*CO_2_; red circles, elevated *p*CO_2_). Shading is in reference to the secondary exposure conditions (light, ambient *p*CO_2_; medium, moderate *p*CO_2_; dark, severe *p*CO_2_). Tertiary exposure conditions, following a 7-day ambient recovery period, are indicated by shape and line type (empty and solid, ambient *p*CO_2_; filled and dashed, moderate *p*CO_2_). Points indicate sampling days for respiration and shell growth measurements and fixed tissues.

### 2.3 Modified reciprocal exposure

#### Secondary Exposure

To begin the secondary exposure, juvenile geoducks (~2,200 geoduck initial *p*CO_2_ treatment^−1^) were rinsed on a 3×10^5^ μm screen to isolate individuals and were divided equally in 36 175-ml plastic cups (*N*=120 animals cup^−1^, *N=6* cups treatment^−1^) each with 50 ml rinsed sand (450-550 μm grain size). Seawater flowed into 250-L head tanks at a rate of 0.6 L min^−1^ and was pumped using submersible pumps to randomly interspersed cups each with a 1 gallon hour^−1^ pressure compensating dripper (Raindrip). Flow rates from dripper manifolds to replicate cups averaged 0.012 liters minute^−1^ (~8 cycles hour^−1^ for 175-ml). Juveniles acclimated under ambient and elevated *p*CO_2_ conditions were subjected to a secondary exposure period (7 days; Fig. 1) in three *p*CO_2_ conditions: ambient (754 ± 15 μatm), moderate (2750 ± 31 μatm), and severe (4940 ± 45 μatm; Table 1).

#### Ambient Recovery

After secondary exposure, *p*CO_2_ addition to head tank seawater ceased and all cups returned to ambient conditions (896 ± 11 μatm, Table 1) for 7 days (Fig. 1).

#### Tertiary Exposure

Replicate cups from the secondary exposure were split (*N*=72 cups) for subsequent tertiary exposure (7 days; Fig. 1) in two conditions: ambient (967 ± 9 μatm) and moderate *p*CO_2_ (3030 ± 23 μatm; Table 1).

Animals were randomly chosen for respiration and growth measurements as described below (*N*=3 geoduck cup^−1^) and fixed in liquid nitrogen (*N*=6 geoduck cup^−1^) every three days and at the start of every treatment transition, cumulatively as days 1, 4, 7 (secondary *p*CO_2_ exposure), 8, 11, 14 (ambient recovery), 15, 18, and 21 (tertiary *p*CO_2_ exposure; Fig. 1). Geoduck were fed *ad libitum* a live mixed-algae diet of *Isocrysis, Tetraselmis, Chaetoceros*, and *Nano* throughout the experiment (4-5×10^4^ cells ml^−1^). Live algae cells were flowed into head tanks during the 21-day modified reciprocal exposure at a semi-continuous rate (2.0 ×10^3^ ml hr^−1^ tank^−1^) with a programmable dosing pump (Jebao DP-4) to target 5×10^4^ live algae cells ml^−1^ in the 175-ml cups. Large algae batch cultures were counted daily via bright-field image-based analysis (Nexcelom T4 Cellometer; (Gurr et al., 2018) to calculate cell density of 2.5×10^4^ live algae cells ml^−1^ in the 250-L head tanks; the closed-bottom cups retained algae to roughly twice the head tank density and algal density was analyzed in three cups via bright field image-based analysis every four days.

### 2.4 Seawater chemistry

Elevated *p*CO_2_ levels in head tanks were controlled with a pH-stat system (Neptune Apex Controller System; Putnam et al., 2016) and gas solenoid valves for a target pH of 7.2 for the elevated and moderate *p*CO_2_ condition and pH of 6.8 for the severe *p*CO_2_ condition (pH in NBS scale). pH and temperature (°C) were measured every 10 seconds by logger probes (Neptune Systems; accuracy: ± 0.01 pH units and ± 0.1°C, resolution: ± 0.1 pH units and ± 0.1°C) positioned in header tanks and trays.

Total alkalinity (TA; μmol kg ^−1^ seawater) of head tank, tray, and cup seawater was sampled in combination with pH (total scale) by handheld probe (Mettler Toledo pH probe; resolution: 1 mV, 0.01 pH; accuracy: ± 1 mV, ± 0.01 pH; Thermo Scientific Orion Star A series A325), salinity (Orion 013010MD Conductivity Cell; range 1 μS/cm - 200 mS/cm; accuracy: ± 0.01 psu), and temperature (Fisherbrand Traceable Platinum Ultra-Accurate Digital Thermometer; resolution; 0.001°C; accuracy: ± 0.05 °C). Quality control for pH data was assessed on each day with Tris standard (Dickson Lab Tris Standard Batch T27). Carbonate chemistry was recorded weekly for each replicate tray during the 110-day acclimation period and daily during the 21-day experiment for three randomized cups representative of each *p*CO_2_ treatment (days 1-7 and 8-15 *N*=9 cups; days 15-21 *N*=6 cups). Additionally, carbonate chemistry of all cups was measured once weekly during each 7-day period (days 1-7 and 8-15, *N*=32 cups; days 15-21, *N*=72 cups). TA was measured using an open-cell titration (SOP 3b; Dickson et al., 2007) with certified HCl titrant (~0.1 mol kg^−1^, ~0.6 mol kg^−1^ NaCl; Dickson Lab, Batches A15 and A16) and TA measurements identified <1% error when compared against certified reference materials (Dickson Lab CO_2_ CRM Batch 180). Seawater chemistry was completed following Guide to Best Practices (Dickson et al. 2007); TA and pH measurements were used to calculate carbonate chemistry, CO_2_, *p*CO_2_, HCO^3-^, CO_3_, and *Ω*_arag_, using the SEACARB package (Gattuso et al., 2018) in R v3.5.1 (R Core Team, 2018).

### 2.5 Respiration rate and shell growth

Respiration rates (oxygen consumption per unit time) were estimated by monitoring oxygen concentration over time using calibrated optical sensor vials (PreSens, SensorVial SV-PSt5-4ml) on a 24-well plate sensor system (Presens SDR SensorDish). Vials contained three individuals per cup filled with 0.2 μm-filtered seawater from the corresponding treatment head tank. Oxygen consumption from microbial activity was accounted for by including 5-6 vials filled only with 0.2 μm-filtered treatment seawater. Respiration rates were measured in an incubator set at 17°C, with the vials and plate sensor system fixed on a rotator for mixing. Oxygen concentration (μg O_2_ L^−1^) was recorded every 15 seconds until concentrations declined to ~50-70% saturation (~20 minutes). Vial seawater volume was measured and clams from each vial were photographed with a size standard (1 mm stage micrometer) to measure shell length (parallel to hinge; mm) using Image J. Respiration rates were calculated using the R package LoLinR with suggested parameters by the package authors (Olito et al., 2017) and following Gurr et al. (2020) with minor adjustments: fixed constants for weighting method (L%) and observations (alpha = 0.4) over the full 20-minute record. Final respiration rates of juvenile geoduck were corrected for blank vial rates and vial seawater volume (μg O_2_ hr^−1^ individual^−1^).

### 2.6 Physiological Assays

Total antioxidant capacity (TAOC), total protein, and ash free dry weight (AFDW; organic biomass) was measured for one animal from each biological tank replicate (*N*=6 animals treatment^−1^) at the end of secondary exposure (36 total animals) and at the end of tertiary exposure (72 total animals). Whole animals were homogenized (Pro Scientific) with 300-500 μl cold 1×PBS and total homogenized volume (μl) was recorded. Homogenates were aliquoted for TAOC and total protein assays and the remaining homogenate was used to measure organic biomass. TAOC was measured in duplicate as the reduction capacity of copper reducing equivalents (CRE) following the Oxiselect™ microplate protocol (STA-360) and standardized to the total protein content of the tissue lysate samples of the same individual (μM CRE mg protein^−1^). Sample aliquots for total protein were solubilized by adding 10 μl 1 M NaOH preceding incubation at 50°C and 800 RPM for 4 hours and neutralized with 0.1 M HCl (pH 7). Total protein of tissue lysate samples was measured using the Pierce Rapid Gold assay with bovine serum albumin following the Pierce™ microplate protocol (A53225). Total protein (mg) was standardized to organic biomass (mg protein mg AFDW) following ignition (4.5 hours at 450°C) subtracted by the dry weight (24 hrs at 75°C) and corrected for total homogenate volume.

### 2.7 Statistical Analysis

Welch’s t-tests for unequal variances were used to analyze the effect of the initial 110-day *p*CO_2_ acclimation (fixed) on respiration rate and shell length prior to the 21-day exposure period. A three-way analysis of variance (ANOVA) was used to analyze the effect of time (fixed) and initial and secondary *p*CO_2_ exposures (fixed) on respiration rate and shell growth for both the secondary exposure and ambient recovery periods (days 1-7 and 8-14, respectively). A four-way ANOVA was used to analyze the effect of time (fixed) and initial, secondary, and tertiary *p*CO_2_ exposures (fixed) on respiration rate and shell growth during the tertiary exposure period (days 15-21). Total antioxidant capacity, total protein, and organic biomass from samples on day 7 and day 21 were analyzed for effects of *p*CO_2_ treatments (fixed) with two-way and three-way ANOVAs, respectively. In all cases, normality assumptions were tested with visual inspection of diagnostic plots (residual vs. fitted and normal Q-Q; (Kozak and Piepho, 2018) and homogeneity of variance was tested with Levene’s test (Brown and Forsythe, 1974). A pairwise Tukey’s *a posteriori* Honestly Significant Difference test was applied to significant model effects. All data analysis was completed using R (v3.5.1; R Core Team, 2018).

## 3. Results

### 3.1 Initial stress acclimation and secondary exposure to hypercapnic seawater

There was no difference in respiration rate after 110 days of *p*CO_2_ acclimation (Table 2; Welch’s t-test; initial, t=-0.602, df=31.725, P=0.5516), however the shell length of geoduck under elevated *p*CO_2_ was significantly larger, by 2.6%, compared to those under ambient treatment (Table 2; Welch’s t-test; initial, t=-4.297, df=2884, P<0.0001). Under subsequent secondary exposure, the initial stress acclimation had a marginal effect on respiration rate (three-way ANOVA; initial, *F*_1,89_=3.409, P=0.068) with 12.4% greater respiration rates in animals under elevated *p*CO_2_ compared to animals under ambient conditions (Table 2). This effect was primarily driven by a marginal interaction effect under secondary *p*CO_2_ treatment (three-way ANOVA; initial×secondary, *F*_2,89_= 2.824, P=0.0647), with 31% greater respiration rates under moderate *p*CO_2_ stress in animals acclimated under elevated *p*CO_2_ (Fig. 2a). Shell length increased significantly with time (three-way ANOVA; time, *F*_2,306_= 3.347, P=0.0365; Tukey HSD, day 7 > day 4, P=0.0236) and independent of *p*CO_2_ treatments (Table 2). Juvenile clams acclimated under elevated *p*CO_2_ on average had significantly greater organic biomass (two-way ANOVA; initial, *F*_1,30_=9.313, P=0.0047) at the end of the secondary exposure period (day 7) with 39% greater mg tissue AFDW individual^−1^ compared to animals reared under ambient conditions (Table 3 and Fig, 3c). There was no significant effect from initial or secondary *p*CO_2_ treatments on total protein or TAOC (Table 3 and Figs. 3a and 3b).

**Table 2.**
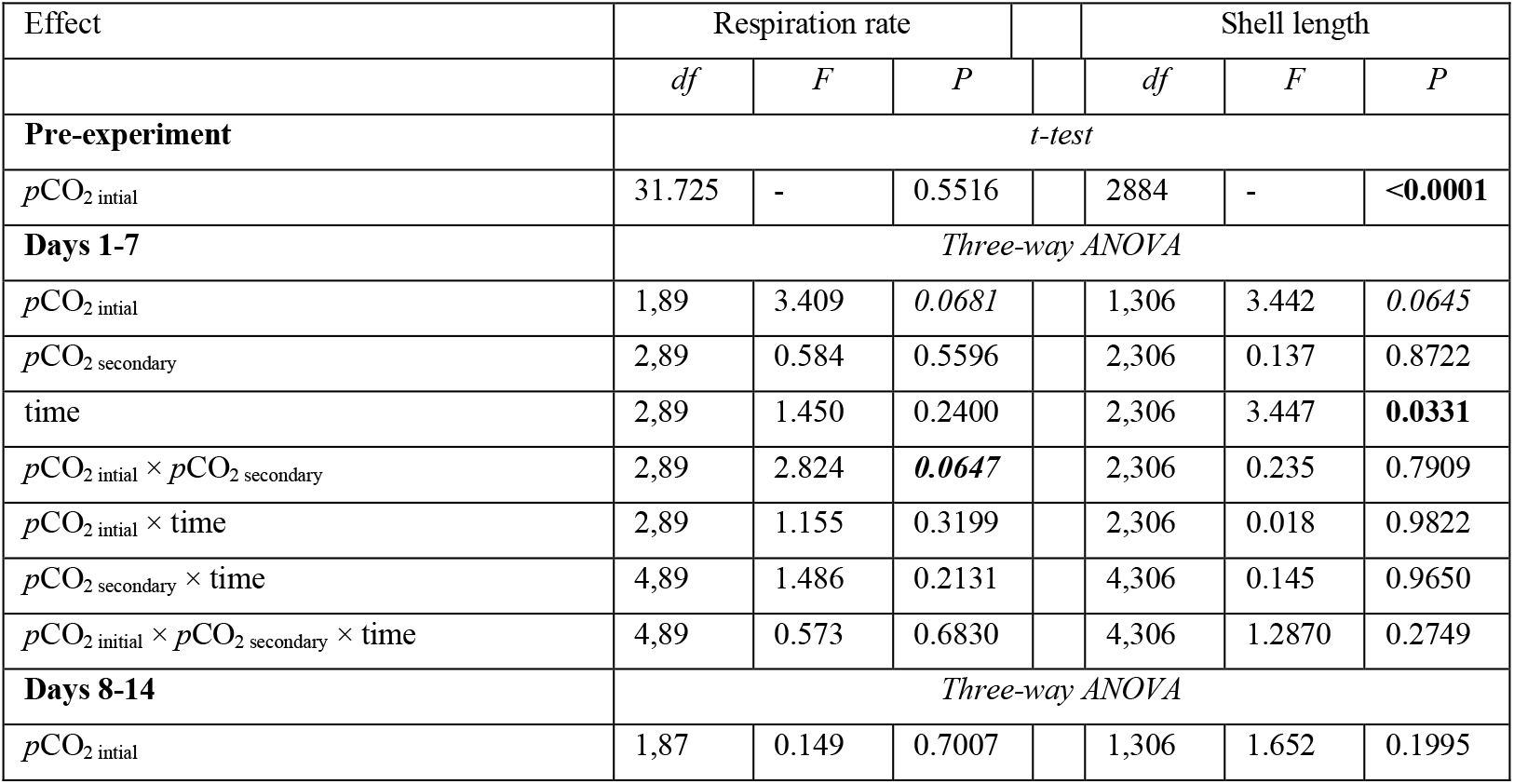
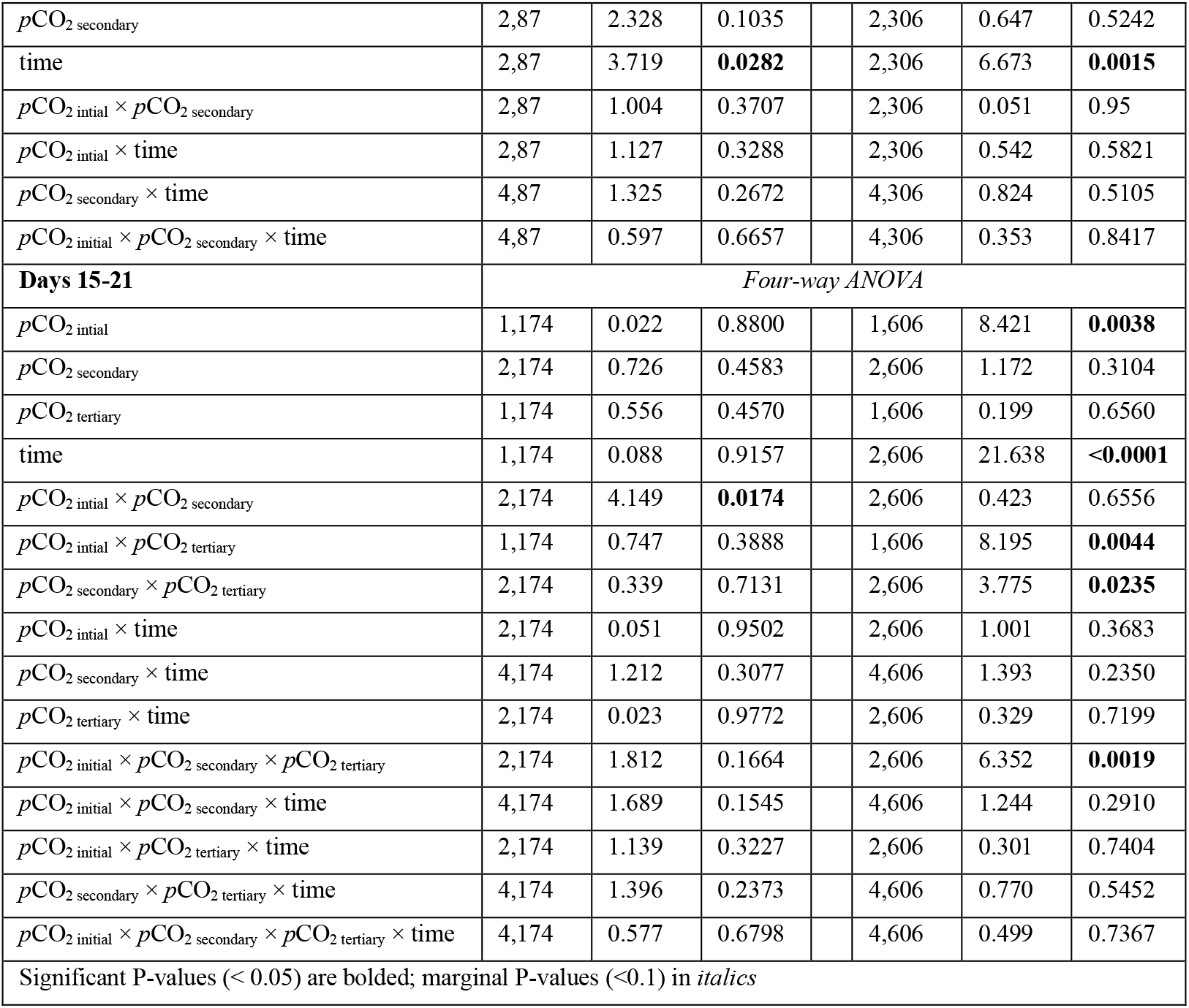
Effects of *p*CO_2_ stress exposures on mean respiration rate and shell growth of *Panopea generosa*. A Welch’s t-test tested for effects of *p*CO_2_ stress acclimation prior to the 21-day experiment. Three-way and four-way ANOVA tests were used under the secondary and tertiary exposure periods and a three-way ANOVA tested for treatment effects during the ambient recovery period. Significant effects are bolded for P < 0.05.

**Fig. 2.**
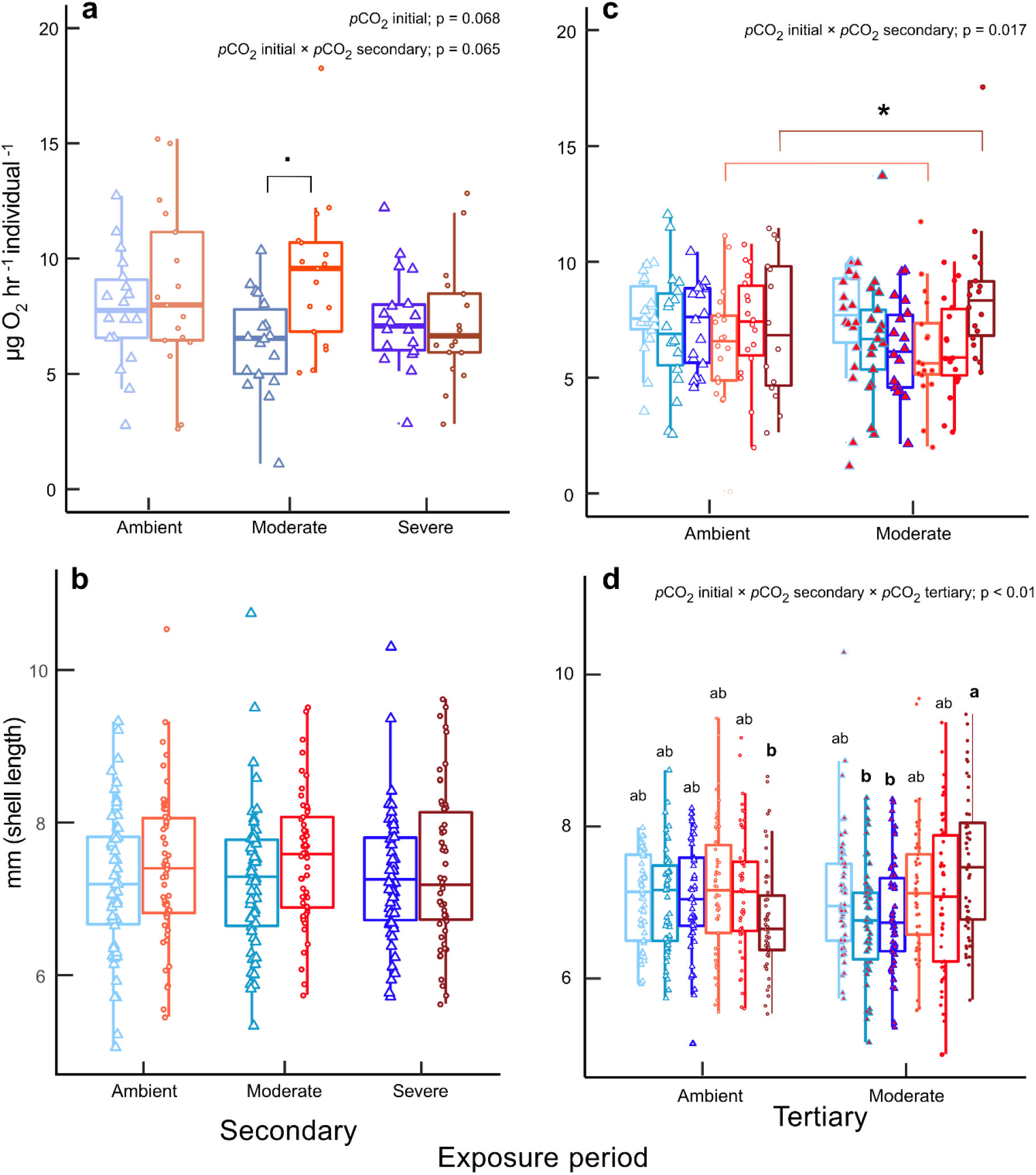
Respiration rate and shell length under secondary (a and b) and tertiary (c and d) exposure periods. Color, shapes, and fill are in reference to Figure 1. Significant *a posteriori* effects are shown as letters or an asterisk.

**Table 3.** Effects of *p*CO_2_ stress exposures on antioxidant capacity, total protein, and organic biomass (AFDW) of *Panopea generosa*. Two-way and three-way ANOVA tests for differences in physiological and cellular status on days 7 and 21 of the 21-day exposure period, respectively. Significant effects are bolded for P < 0.05.

| Effect | Antioxidant capacity |  |  | Total protein |  |  | AFDW |  |  |
| --- | --- | --- | --- | --- | --- | --- | --- | --- | --- |
|  | *df* | *F* | *P* | *df* | *F* | *P* | *df* | *F* | *P* |
| **DAY 7** | *Two-way ANOVA* |  |  |  |  |  |  |  |  |
| $p\text{CO}_2$ initial | 1,30 | 0.005 | 0.942 | 1,30 | 0.003 | 0.959 | 1,30 | 9.313 | **0.0047** |
| $p\text{CO}_2$ secondary | 2,30 | 0.143 | 0.867 | 2,30 | 0.866 | 0.431 | 2,30 | 2.536 | 0.096 |
| $p\text{CO}_2$ initial × $p\text{CO}_2$ secondary | 2,30 | 1.007 | 0.377 | 2,30 | 2.136 | 0.136 | 2,30 | 0.158 | 0.8546 |
| **DAY 21** | *Three-way ANOVA* |  |  |  |  |  |  |  |  |
| $p\text{CO}_2$ initial | 1,56 | 8.069 | **0.0063** | 1,56 | 2.365 | 0.13 | 1,56 | 12.899 | **<0.001** |
| $p\text{CO}_2$ secondary | 2,56 | 0.164 | 0.849 | 2,56 | 0.625 | 0.539 | 2,56 | 1.578 | 0.2153 |
| $p\text{CO}_2$ tertiary | 1,56 | 2.161 | 0.1471 | 1,56 | 1.272 | 0.264 | 1,56 | 3.298 | *0.0747* |
| $p\text{CO}_2$ initial × $p\text{CO}_2$ secondary | 2,56 | 1.43 | 0.248 | 2,56 | 1.423 | 0.25 | 2,56 | 1.756 | 0.1822 |
| $p\text{CO}_2$ initial × $p\text{CO}_2$ tertiary | 1,56 | 0.678 | 0.4136 | 1,56 | 2.25 | 0.139 | 1,56 | 0.453 | 0.5036 |
| $p\text{CO}_2$ secondary × $p\text{CO}_2$ tertiary | 2,56 | 0.752 | 0.476 | 2,56 | 0.225 | 0.8 | 2,56 | 0.166 | 0.8906 |
| $p\text{CO}_2$ initial × $p\text{CO}_2$ secondary × $p\text{CO}_2$ tertiary | 2,56 | 0.141 | 0.8688 | 2,56 | 0.26 | 0.772 | 2,56 | 0.181 | 0.8353 |
| Significant P-values (< 0.05) are bolded; marginal P-values (<0.1) in *italics* |  |  |  |  |  |  |  |  |  |

**Fig. 3.**
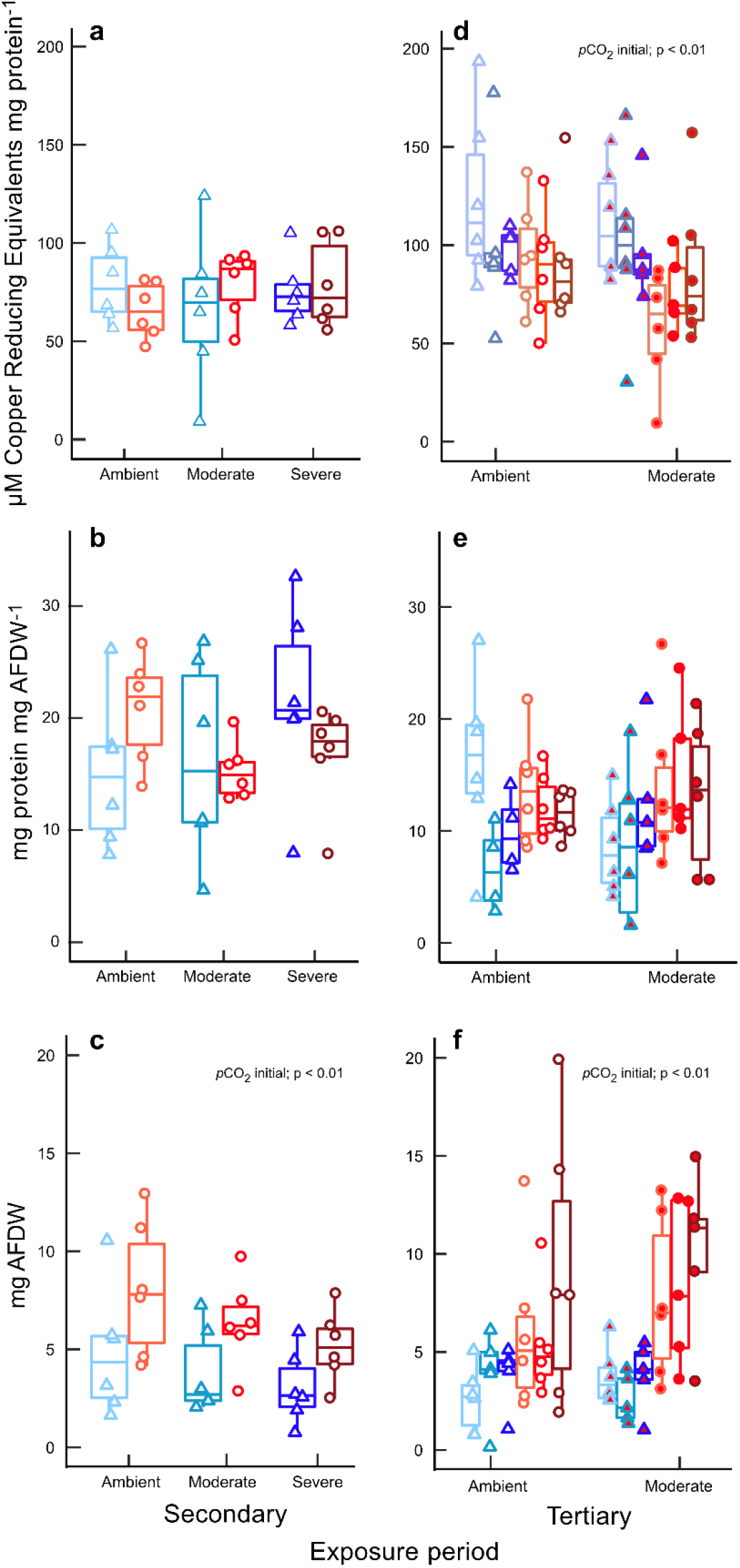
Antioxidant response and physiology of fixed animals at the end of secondary (a-c; day 14) and tertiary treatments (d-f; day 21). Color, shapes, and fill are in reference to Figure 1.

### 3.2 Ambient common garden recovery under normocapnic conditions

During ambient recovery, respiration rate was not significantly affected by initial or secondary *p*CO_2_ treatments, but was significantly affected by time (Table 2; three-way ANOVA; time, *F*_2,87_=3.719, P=0.0282) with an increase in respiration rates over the 7-day period (Tukey HSD; day 14 > day 8; P = 0.0254). Similarly, shell growth was not significantly affected by initial or secondary *p*CO_2_ treatment, but showed a significant increase with time (Table 2; three-way ANOVA; time, *F*_2,306_=6.643, P=0.0015; Tukey HSD; day 14 > day 8, P=0.0.013; Tukey HSD, day 11 > day 8, P=0.001).

### 3.3 Tertiary exposure to hypercapnic seawater

The interaction of initial and secondary *p*CO_2_ treatments had a significant effect on respiration rate under tertiary exposure (Table 2; four-way ANOVA; initial×secondary, *F*_2,174_= 4.149, P=0.0174), with this interaction primarily driven by a 20.4% greater respiration rate in *p*CO_2_ stress-acclimated animals exposed to severe and ambient *p*CO_2_ during the secondary period (Fig. 2c), although the post-hoc test was only marginally significant (Tukey HSD; moderate×severe > moderate×ambient, P=0.0685). Shell growth was affected by an interaction between initial, secondary, and tertiary *p*CO_2_ treatments (Table 2 and Fig. 2d; four-way ANOVA; initial×secondary×tertiary, *F*_2,606_=6.352, P=0.0019). Pairwise differences of the three-way treatment interaction showed 9.3% greater mean shell size by acclimated animals with secondary and tertiary exposure to severe and moderate *p*CO_2_, respectively (Fig. 2d). At the end of the tertiary exposure period (day 21), initial stress acclimation under elevated *p*CO_2_ increased organic biomass (Table 3; three-way ANOVA; initial, *F*_1,56_=12.899, P<0.001) and tertiary exposure had a marginal increase (three-way ANOVA; tertiary, *F*_1,56_=3.298, P=0.0604) with 51% and 28% greater organic biomass under stress treatments relative to ambient controls (Fig. 3f). There was a significant effect of initial stress acclimation on antioxidant activity (Table 3; three-way ANOVA; initial, *F*_1,56_=8.069, P=0.0063) with 22% greater μM CRE mg protein^−1^ by clams reared under ambient *p*CO_2_ (Fig. 3d); there was no effect of *p*CO_2_ treatment or two-way and three-way interactions of *p*CO_2_ treatments on total protein (Table 3 and Fig. 3e).

## 4. Discussion

In the present study we evaluated the effects of postlarval stress acclimation and subsequent exposures to elevated *p*CO_2_ on the physiological and oxidative stress response in juvenile geoduck. Our findings suggest moderate hypercapnic conditions during postlarval development improves metrics of physiological performance and CSR. This novel investigation of adaptive stress acclimation demonstrates a high tolerance to *p*CO_2_ regimes (~2500-5000 μatm) and plasticity of bioenergetic and subcellular responses beneficial for later performance in *Panopea generosa*.

### 4.1 Stress-intensity- and life-stage-dependent effects

Survival under long-term stress exposure and positive physiological responses of acclimated animals under ‘moderate’ (~2900 μatm *p*CO_2_ 0.4 *Ω*) and ‘severe’ (~4800 μatm *p*CO_2_ 0.2 *Ω*) reciprocal exposures highlights the resilience of *Panopea generosa* to OA and suggests that stress acclimation (days) can induce beneficial effects during postlarval-juvenile development. Specifically, clams repeatedly exposed to the greatest intensity of stress (moderate×severe×moderate) had both greater respiration rates and shell size (Table 2; Fig. 2). Further, stress-acclimated individuals had greater organic biomass and lower amounts of antioxidant proteins relative to ambient controls (Figs. 3c, 3d, and 3f), suggesting optimized tissue accretion and energy partitioning, coupled with decreased costs for cytoprotection. Previous studies describe metabolic compensation and regulation of CSR during hypercapnia as attributes of a well-adapted stress response to control acid-base status and normal development/metamorphosis (Walsh and Louise Milligan, 1989; Dineshram et al., 2015). Indeed, prior work on juvenile *P. generosa* also demonstrates positive acclimatory carryover effects, with increased shell length and metabolic rate after repeat exposures to hypercapnic and undersaturated conditions with respect to aragonite (Gurr et al., 2020). Contrary to these findings, similar *p*CO_2_ and *Ω* levels decrease metabolic rate and scope for growth in the mussel *Mytilus chilensis* (Navarro et al., 2013), cause a three-fold increase in mortality rate in juvenile hard clam *Mercenaria mercenaria* (Green et al., 2009), and alter metamorphosis and juvenile burrowing behavior in *Panopea japonica* (Huo et al., 2019). Thus,*p*CO_2_ tolerance limitations are likely species-specific, as well as life stage, duration, and stress-intensity specific.

*p*CO_2_-induced phenotypic variation over postlarval-juvenile development observed in this study suggests postlarval stages may be optimal for stress acclimation. Beneficial carryover effects herein are also corroborated with compensatory physiology and differential DNA methylation of juvenile *P. generosa* in other studies (Putnam et al., 2017; Gurr et al., 2020). In contrast, OA can have deleterious effects on growth/development, settlement, and proteomic composition of larval *P. generosa* (Timmins-Schiffman et al., 2020), further emphasizing life-stage dependence of *p*CO_2_ stress exposure. Mollusc larvae are widely established to have enhanced susceptibility to OA with impacts on shell growth and developmental transition (Kurihara et al., 2007; Kapsenberg et al., 2018). For example, larval exposure to elevated *p*CO_2_ leads to persistent negative effects (i.e. reduced shell growth and development) in Pacific oyster *Crassostreas gigas*, Olympia oyster *Ostrea lurida*, and bay scallop *Argopecten irradians* (Barton et al., 2012; Hettinger et al., 2012; White et al., 2013). Beneficial responses to OA are also possible, especially in longer term and carryover-effect studies (Parker et al., 2015). For example, elevated *p*CO_2_ during gametogenesis in the Chilean mussel *Mytilus chilensis* and Sydney rock oyster *Saccostrea glomerata* increases the size of larval stages in progeny (Parker et al., 2012; Diaz et al., 2018). Future comparative studies should test stress responses of well-defined inter- and subtidal molluscs to determine if pre-conceived tolerance limitations are affected by repeated dose-responses post-settlement.

Our observation of beneficial effects in stress-acclimated clams suggest an adaptive resilience of *P. generosa* to hypercapnic conditions relevant to postlarval-juvenile development in both natural and aquaculture systems. *P. generosa* are likely capable of adaptive resilience particularly during this life stage because *p*CO_2_ and *Ω* gradients naturally occur alongside the dramatic developmental transition from free-swimming larvae to sessile benthic juveniles. Habitat within the native range of *P. generosa* exhibits elevated *p*CO_2_ and aragonite undersaturation with episodic/seasonal variation (surface water *Ω*<1 in winter months, Dabob Bay in Hood Canal, WA; Fassbender et al., 2018) and geographical (>2400 μatm and *Ω*<0.4 in Hood Canal, WA; Feely et al., 2010) and vertical heterogeneity (Reum et al., 2014) comparable to gradients within sub-surface sediments (*Ω* 0.4-0.6; Green et al., 2009). Meseck et al. (2018) found *in-situ* porewater pH increases model predictability of bivalve community composition, *Mya arenaria* and *Nucula* spp, suggesting sediment acidification and carbonate chemistry influence bivalve settlement. Our experimental timing and findings overlaid on common geoduck aquaculture practices suggest postlarval ‘settlement’ as an ecologically-relevant window to investigate the adaptive capacity for stress acclimatization.

### 4.2 Oxidative status and repeated stress encounters

Our results herein demonstrate activation of phenotypic variation after repeated stress encounters suggesting postlarval stress acclimation may have a critical role in subsequent stress response. A growing body of research supporting conditioning-hormesis challenges the paradigm of primarily deleterious effects of stress exposure (Calabrese et al., 2007; Costantini et al., 2012; Gurr et al., 2020). Here we posit that hormetic priming could be occurring through moderate oxidative stress on protein homeostasis (patterns of differential expression), cellular signaling, and mitochondrially-mediated responses. For example, Constantini et al. (2012) found early-life exposure of the zebra finch *Taeniopygia guttata* to thermal-induced oxidative damage decreased oxidative stress under subsequent thermal exposure during adulthood. This low-dose stimulatory effect of oxidative stress is well-characterized (i.e. under calorie restriction, hypoxia, and exercise; Ristow and Schmeisser, 2014) for a wide range of taxa (Costantini et al., 2012; Visser et al., 2018; Zhang et al., 2018), but remains poorly understood in response to OA conditions.

In *Panopea generosa* in this study, compensatory antioxidant synthesis reduced performance in absence of prior stress acclimation suggesting an adaptive role of oxidative stress and conditioning-hormesis. pH- and CO_2_-induced oxidative stress and antioxidant response are of growing interest to characterize stress resilience in bivalves (Tomanek et al., 2011; Matoo et al., 2013). Oxidative stress is intensified by acidosis (low intracellular pH) and hypercapnia either indirectly from low pH on ferrous iron enhancing the Fenton reaction (producing hydroxyl radicals), or a direct interaction of intracellular hypercapnia on free radical formation (Tomanek et al., 2011). pH- or CO_2_-induced stress and mediative responses are species and stress-dependent (i.e. frequency, duration, and magnitude of stress exposures). Under short and prolonged exposure to subtle hypercapnic seawater (2-15 weeks; ~800 μatm, 7.9 pH, and 3 *Ω*), the Eastern oyster *Crassostrea virginica* and hard clam *M. mercenaria* differ in initial antioxidant production suggesting that CSR can determine interspecies vulnerability to hypercapnic seawater (Matoo et al., 2013). In more pronounced hypercapnic and aragonite undersaturated conditions, *C. virginica* upregulates antioxidant proteins (14 days; ~3500 μatm and 7.5 pH; Tomanek et al., 2011) and a similar response is also seen in the Yesso scallop *Patinopecten yessoensis* (duration = 14 days; ~2200 μatm, 7.5 pH, and 0.7 *Ω;* Liao et al., 2019) and mussel *Mytilus coruscus* (duration = 45 days; ~2800 μatm, 7.3 pH, and 0.5 *Ω*; Huang et al., 2018). Alternatively, a seemingly ‘preparatory’ energetic cost of antioxidant synthesis is found in a variety of taxa (Hermes-Lima and Zenteno-Savín, 2002; Ivanina and Sokolova, 2016) and suggests adaptive energy reallocation to scavenge ROS formed by aerobic recovery when a stressor is lifted.

Intermittent oxidative stress may have evolutionary importance in stress resilience of long-lived marine bivalves. The ocean quahog *Arctica islandica* is the oldest known non-colonial animal; their substantial longevity is hypothesized to be driven by intermittent metabolic-quiescence (dormancy when burrowed) demanding resilience to ROS overproduction (oxidative bursts) and resistance to cell death upon subsequent aerobic-recovery (Abele et al., 2008). Interestingly, *A. islandica* have a particularly low peroxidation-sensitive lipids (Munro and Blier, 2012) and high baseline antioxidant capacity throughout their lifespan suggesting an adaptive resilience to oxidative damage (Abele et al., 2008). Whereas intertidal molluscs (i.e. *Mytilus sp., Crassostrea sp*., and *Argopecten sp*.) have energetic deficits under dynamic environmental stress (i.e. elevated metabolic demand, ROS production, and antioxidant response; Kelley et al., 2017; Liao et al., 2019), *A. islandica* continues ventilation without anaerobic transition or overproduction of free radicals (Strahl et al., 2011). Thus, lower antioxidant production by stress-conditioned *P. generosa* could suggest adaptive subcellular mechanism(s) common across long-lived bivalves to maintain redox status under frequent or intermittent stress exposures. Altogether, the contrasting response of lower antioxidant proteins in stress-acclimated *P. generosa* suggests beneficial effects of prior exposure to hypercapnic seawater, which highlights the need for a mechanistic understanding of the role of oxidative stress in this process.

### 4.3 Mitochondrial and molecular mechanisms of moderate stress acclimation

Based on the effects of stress acclimation on antioxidant capacity and performance of *Panopea generosa* in this study, we describe two possible subcellular mechanisms underpinning stress-intensity-dependent effects on mitochondrial pathways and energy partitioning: 1) stress-induced attenuation of mitochondrial function and 2) adaptive mitochondrial shift under moderate stress (Fig. 4).

**Fig. 4.**
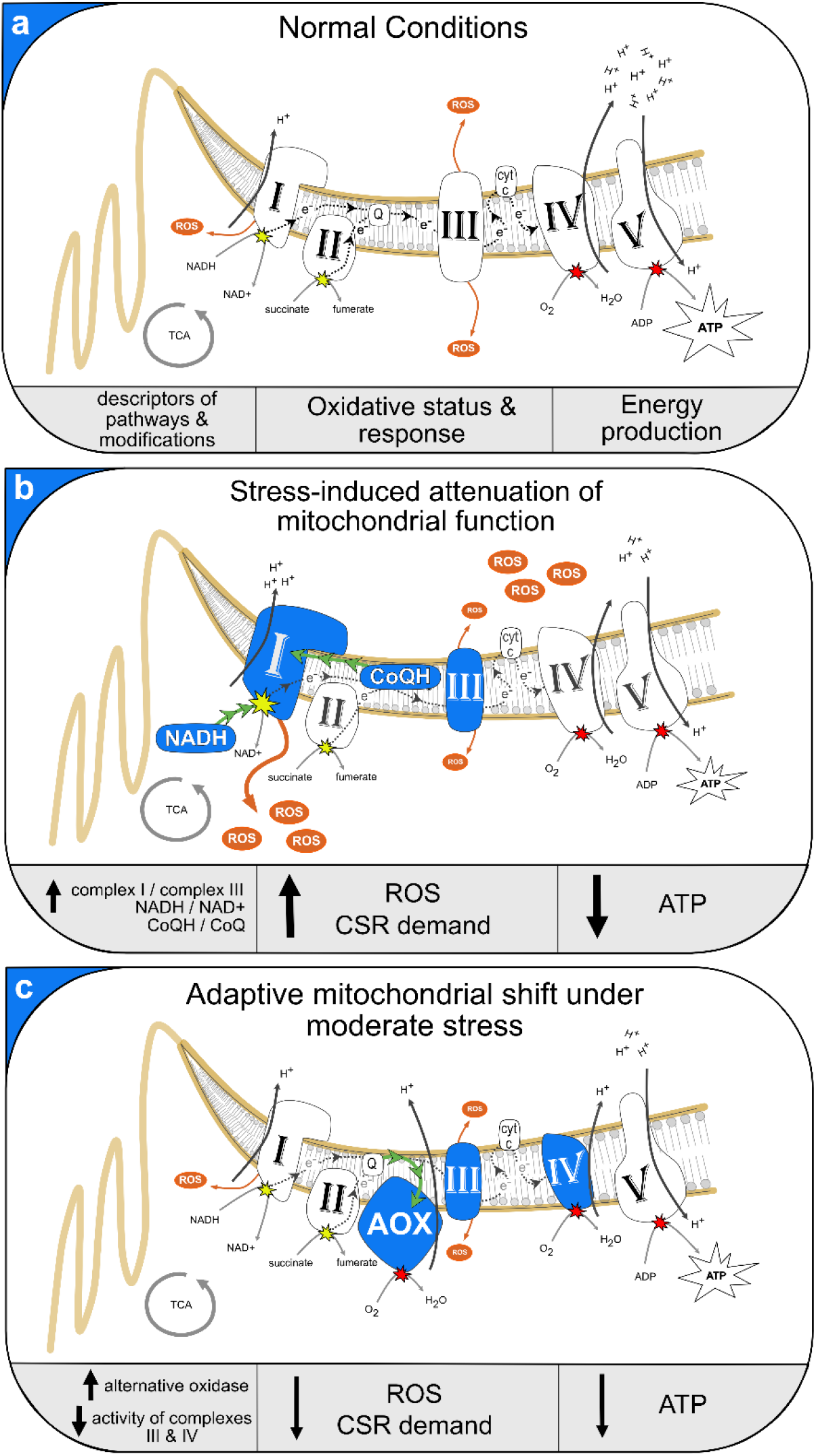
Hypothesized model of stress acclimation through mitochondrial pathways: Diagram of inner mitochondrial membrane under normal conditions (a), dysfunction (b; i.e. reverse electrician transport), and the AOX pathway (c). Solid blue shapes highlight descriptors of modified pathways and green arrows display shifts in the direction of electrons (b and c). Legends with solid black arrows display the direction of changes relative to the mETC under normal conditions.

#### 1) Stress-induced attenuation of mitochondrial function

Individuals initially exposed to ambient *p*CO_2_ increased antioxidant proteins and decreased growth relative to stress-acclimated clams suggesting a greater demand for CSR leading to negative performance. Hypercapnia and acidosis can directly and indirectly drive increased formation of free radicals (Murphy, 2009; Tomanek et al., 2011) and elicit changes in the activity (Lambert and Brand, 2004) and expression of mitochondrial complexes (Dineshram et al., 2012; Dineshram et al., 2015), which altogether suggests diminished mitochondrial ATPs under environmental stress (Murphy 2009). Under normal aerobic respiration (Fig. 4a), complexes I (NADH dehydrogenase) and III (cytochrome *c* oxidoreductase) are dominant sources of mitochondrial ROS production by the electron transport chain (mETC). In contrast, mitochondrial dysfunction or alterations of mETC occur under environmental stress in which high pools of NADH in the mitochondrial matrix and reduced coenzyme Q shift electron transport (reverse electron transport or RET) and proton gradient control to complex I, eliciting a deficit for ATPs and enhanced ROS production and CSR demand (Murphy, 2009); Fig 4b). Further, increased protonmotive force and pH gradient of the inner mitochondrial membrane increases RET and ROS production (Lambert and Brand, 2004; Miwa and Brand, 2003) suggesting a mechanism of pH- and CO_2_-induced mitochondrial dysfunction. Altogether, the upregulation of antioxidant proteins and reduced growth in unconditioned *P*. *generosa* are likely outcomes of acclimation-dependent attenuation of mitochondrial function (Fig. 4b). In addition, uncoupling proteins (UCPs) are activated under oxidative damage to mediate mitochondrial stress although further decreasing the proton gradient and ATP production (Slocinska et al., 2016).

#### 2) Adaptive mitochondrial shift under moderate stress

Alternative oxidase (AOX) is a regulatory protein of the inner mitochondrial membrane that receives electrons from ubiquinol to reduce oxygen and produce ATP without involving the cytochrome *c* mETC pathway (Figs. 4a and 4c). There is growing interest in AOX as a compensatory response that permits ATP synthesis and reduces ROS production during stress exposure (Tschischka et al., 2000; Sussarellu et al., 2013; Yusseppone et al., 2018). Although previously theorized as an exclusive mechanism in plant mitochondria (Vanlerberghe, 2013), molecular tools have helped define an evolutionary diversification of AOX in higher order animalia (McDonald and Vanlerberghe, 2006) with a particular prevalence in marine invertebrates and bivalve genomes (McDonald et al., 2009). AOX permits oxidative phosphorylation when complexes III and IV are inhibited (i.e. cyanide and sulphide inhibition), or when ROS is overproduced. A pH decline within the inner mitochondrial membrane can also activate the AOX pathway (Lima-Júnior et al., 2000). Although less efficient, with one-third of the membrane potential relative to cytochrome *c* mETC pathway (Millenaar and Lambers, 2003), AOX activity protects against free radical formation under abiotic and biotic disturbances (McDonald et al., 2009). Upregulated AOX coupled with decreased antioxidant protein synthesis under thermal stress suggests regulatory control of redox status, cellular signaling, and downstream energy allocation (Maxwell et al., 1999) are possible using this pathway. For example, *C. gigas* upregulates AOX in response to hypoxia-reoxygenation suggesting anticipatory dissipation of the proton gradient to avoid overproduction of free radicals upon reoxygenation (Sussarellu et al., 2013). An AOX-mediated response corroborates our hypothesis of a mechanism for hormesis that may outweigh energetic costs of the reduced ATP efficiency from this AOX pathway. A BLAST search identified AOX, mitochondrial carrier proteins (UCPs), and conserved mETC complexes within an annotated Pacific geoduck draft genome (Roberts et al., 2020). qPCR and metabolomics are planned to investigate these proposed mechanisms of hypercapnic-induced modification of mitochondrial function and link complexes I and III, AOX, reduced cofactors, UCPs, and ATP under acute and repeated stress encounters.

## 5. Conclusion

Postlarval acclimation under moderate hypercapnia can elicit beneficial phenotypes under subsequent stress encounters. This acclimatory capacity is likely contingent on stress-intensity (i.e. magnitude, duration, frequency of stress periods) and timing during postlarval settlement and juvenile development. Thus, investigations of marine species responses to climate change should consider adaptive dose-dependent regulation and effects post-acclimation. Further, alternative mitochondrial pathways can build an understanding of mechanisms underpinning hormesis to provide additional ‘climate-proofing’ strategies in aquaculture and conservation of goods and services in the Anthropocene.

## Acknowledgments

We thank the Jamestown S’Klallam Tribe and Kurt Grinnell for providing the animals and facilities for this research. We also thank management staff and technicians at the Jamestown Point Whitney Shellfish Hatchery, Matt Henderson, Josh Valley and Clara Duncan, for their assistance. We also thank Emma Strand for analytical support.

## Author contributions

S.J.G., B.V., S.B.R. and H.M.P. designed the experiments, S.J.G. conducted the experiments and analyzed data, S.J.G., S.A.T, B.V., S.B.R. and H.M.P. drafted, revised, read and approved the final version of the manuscript for publication.

## Data availability

All raw data and statistical code are openly available as a Zenodo repository: http://doi.org/10.5281/zenodo.3903019 (Gurr et al. 2020).

